# CRIS: A Centralized Resource for High-Quality RNA Structure and Interaction Data in the AI Era

**DOI:** 10.1101/2025.05.05.652292

**Authors:** Wilson H. Lee, Cynthia Dharmawan, Kongpan Li, Jianhui Bai, Pratham Solanki, Aman Sharma, Minjie Zhang, Zhipeng Lu

**Author notes:** Authors contributed equally to this work (Wilson H. Lee, Cynthia Dharmawan).

## Abstract

As interest in RNA-based therapeutics expands, there is a growing demand for RNA structure elucidation and RNA-RNA interactions in both academic and clinical settings. Despite rapid advances in methods for RNA structure determination, the field faces persistent challenges in data reproducibility, quality control, and accessibility, largely due to inconsistencies in data processing and analysis workflows. Concurrently, methodological improvements have generated increasingly complex datasets, which necessitate a standardized framework. Here, we present the Crosslinking-based RNA Interactomes and Structuromes (CRIS) database, a comprehensive resource designed to address these limitations. Among existing experimental and computational approaches for RNA structure characterization, crosslinking-based technologies offer superior reliability, high throughput, and high resolution. CRIS provides rigorously curated datasets, standardized workflows, and user-friendly tools, together with built-in quality metrics and detailed visualization guidance to ensure reproducibility and transparency while pairing seamlessly with existing experimental pipelines. By delivering high-complexity RNA datasets alongside accessible computational tools, CRIS serves as a standardized reference for both new and existing data, facilitating investigation through comparative analyses and providing a training resource for deep learning-based computational exploration. This will enable integration into machine learning workflows for large scale, novel RNA structure discovery.

## INTRODUCTION

Once considerably solely as an intermediary between DNA and protein, RNA is now recognized as a key regulator of cellular processes across diverse organisms and viruses (1–3). Recent advancements in methods development have unveiled complex structure-dependent functions and networks of RNA-RNA and RNA-protein interactions that drive processes like gene regulation, splicing, epigenetic modifications, positioning RNA as a critical influencer of cellular function and disease (2–9).

The therapeutic potential of targeting RNA is undeniable (10–12). Successful development of mRNA vaccines for SARS-CoV-2 and antisense oligonucleotides (ASOs) for genetic disorders like spinal muscular atrophy underscore the transformative effects of RNA therapeutics (10,13–17). Insights into viral RNA structures have also informed antiviral strategies, including targeting internal ribosome entry sites (IRES) in pathogens like Hepatitis C virus or poliovirus, and the 3’-UTR in West Nile virus to disrupt viral replication (18–20). It is important to note that these advances relied on precise RNA structure insights; for instance, the SARS-CoV-2 vaccine efficacy is dependent on 5’-UTR stability and required targeting knowledge of splice-site accessibility (21). Despite these advances, characterizing RNA structures *in vivo* remains a significant challenge (1,22–24). While traditional high-resolution methods like X-ray crystallography, cryo-electron microscopy, and NMR provide precise structural insights, they are limited by technical constraints, sample requirements, and potential artifacts (23,25–29). High-throughput chemical probing techniques, leverage the reactivity of RNA nucleotides to map the potential interactions, but they still lack discreet evidence of structure formation (30–33). Methods developed in our group, such as PARIS and SHARC, combine crosslinking chemistry and proximity ligation with next-generation sequencing to capture RNA interactions in their native context, simultaneously providing critical data in secondary and tertiary structure mapping (34–36). Thus, the rapid generation of data through these methods highlights the need for rigorous computational analysis to ensure reproducibility, quality control, and accurate biological interpretation (24,27,37–39).

Due to the complexity of RNA structural data obtained through various methodologies, there is a critical need for standardized, high-quality pipelines that support experimental reproducibility and scalable computational modeling (40–44). Other current databases have limited data curation methods, built-in quality metrics, and integration capabilities for machine learning frameworks, restricting their utility in predictive modeling, large-scale data mining, and novel RNA structure discoveries (40,45).

The Crosslinking-based RNA Interactomes and Structuromes (CRIS) database thus integrates comprehensive datasets with robust analysis steps and visualization tools, providing a foundation for accelerated discoveries in RNA biology, therapeutic development, and the broader field of translational research. CRIS focuses on rigorous, experimentally derived dataset-specific analysis, providing standardized processed data and direct links to files and instructions when appropriate. It does not attempt unreliable global meta-analyses across heterogeneous RNA types, ensuring high-confidence structural and interaction information.

Given the expansive implications of RNA in disease states, precise structural understanding enabled by comprehensive datasets will guide the rational design of therapeutically functional RNAs. For instance, structure data can inform RNA aptamer design to specifically bind RNA and protein targets (46–48). Understanding how fundamental structural elements govern folding or splicing mechanisms not only accelerates the development of novel RNA-based therapeutics but also deepens our insight into how specific RNA motifs dictate binding, regulation and functional outcomes. This knowledge will replace trial- and error approaches as rule-based, predictive frameworks will be developed based on *in vivo* structure data. Coupled with built-in quality metrics, user-friendly guides, and seamless integration with widely used bioinformatics tools, CRIS empowers researchers to explore RNA structure and function with unprecedented precision and scalability. In doing so, CRIS establishes a new standard for RNA informatics, fostering innovation and transformative breakthroughs across biological and medical sciences.

## DATABASE CONTENT

### CRIS Database Overview

The CRIS database represents a significant advancement in RNA structure research by harmonizing crosslink-ligate RNA sequencing data captured by a variety of crosslinkers across *Homo sapiens, Mus musculus*, and various viral genomes CRIS utilizes rigorous quality control (QC) and consistent data processing, addressing a key gap in existing databases where inconsistent or unavailable QC metrics hinder reliable cross-study comparisons. By providing processed data files (i.e. aligned, crosslink classified, duplex groups) ready for further analyses of individual RNAs or comparisons, CRIS enables a precise exploration of RNA structures and interactions.

The web interface caters to users across various experience levels, accommodating both quick browsing of data statistics for entry-level users as well as flexible command-line scripts for advanced users. The landing page provides an overview of CRIS and its capabilities, and we have included more detailed sections to allow for efficient workflow navigation. Specifically, we highlight the glossary page that details terminology we standardized to aid understanding and maximize adoptability among the user base, a statistics page that summarizes experimental data metrics, enhancing transparency, and a tools page that includes a quick start guide to reproduce the same analysis. This tools page is further sectioned into step-by-step instructions (with example command-line code), enabling users to move from data download to publication-ready results with minimal complexity.

Data rich intermediary files detailing RNA crosslinking (XLRNA) configurations (e.g., gap1, gapm files) are provided through their respective download pages for SHARC and PARIS data, but CRIS does not store any raw sequencing data as those files are already accessible through NCBI GEO or ENA. As such, CRIS is not a service provider; it does not host source files or provide computational power for data analysis. Rather, its purpose is a collection of the rigorously curated, highest quality datasets to empower researchers to conduct their own deeper analyses with confidence. CRIS prioritizes reproducible workflows and detailed documentation. The interface design includes intuitive download options, tutorials for data visualization, and quick statistical summaries table (e.g., read counts, segment lengths) that can be used to compare datasets. Users can follow the data from start-to-finish, guided stepwise from data collection to visualization without a prior coding background. Note that the processed data does not serve as a platform for any global meta-analyses because this approach often leads to faulty comparisons.

### Data Curation

CRIS includes a broad range of crosslink-ligation sequencing datasets, including PARIS1/2 (GSE74353, GSE149493), SPLASH (PRJNA318958), LIGR-seq (GSE80167), COMRADES (E-MTAB-6427), SHARC (GSE167812), hiCLIP (E-MTAB-2937, E-MTAB-2940, E-MTAB-2941), RIC/cRIC-seq (GSE127188, GSE210583), as well as extensive miRNA data, selected for their variety in crosslinking technology and strengths in capturing RNA structures and interactions (34,35,49–58). These datasets cover diverse biological contexts, from various cell lines and model organisms to virus-host interactions, enhancing the utility of CRIS for comparative analyses across species and experimental conditions. We reprocessed all public datasets (e.g., PARIS, RIC/cRIC-seq) from raw FASTQ files using the standardized pipeline (59). To ensure consistency, files were renamed with a structured convention: [Species]-*[CellLine]-* [Crosslinker]-*[LigationTime]-*[RNaseRTime] (e.g., HS-HEK293T_Amo-0.5_T4-24h_exo-0h). For example, “HS-HEK293T_Amo-0.5_T4-24h_exo-0h” refers to data generated using samples of homo sapiens origin, HEK293T cell line, amotosalen crosslinker at 0.5 mg/mL, proximally ligated for 24 hours with T4 RNA ligase, and not treated with RNase R exonuclease.

XLRNA data generated internally from PARIS and SHARC experiments were preprocessed for adapter removal, demultiplexes, and PCR duplicate removal by a wrapper script. This pipeline utilizes scripts from the PARIS/SHARC computational pipeline for streamlined preprocessing; processing tools used in CRIS (e.g. Trimmomatic, STAR) were selected for flexibility in parameter customization (**Figure 1**). This allows for tailored analyses of crosslinking datasets, an area where commercially available software packages have processing limitations. All data available on CRIS were aligned to a modified, masked reference genome using optimized STAR parameters prior to duplex group (DG) assembly using CRSSANT scripts (37). This approach ensures high-precision mapping and standardized classification of RNA structures and interactions.

**Figure 1.**
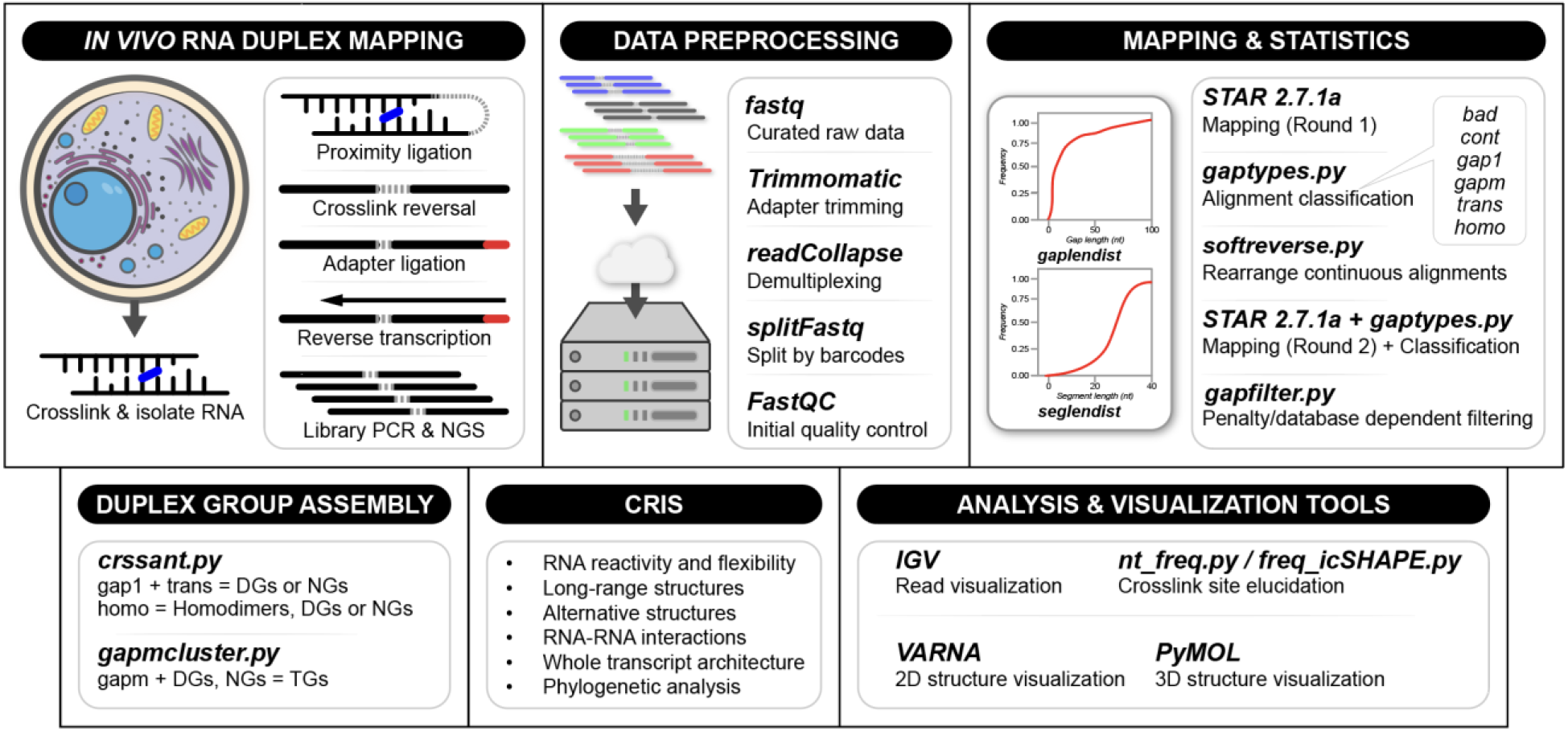
Workflow overview for *in vivo* RNA duplex mapping and CRIS-based analysis. The pipeline begins with *in vivo* RNA crosslinking, followed by RNA isolation and preparation of cDNA libraries for high-throughput sequencing. Raw sequencing data undergo preprocessing steps including adapter trimming, barcode splitting, and quality control via the wrapper script. Reas are then aligned to a reference genome using a two-round mapping and classification strategy, with intermediate analysis of alignment types and structural gap features. Duplex group assembly is performed using *crssant.py* and *gapmcluster.py*, enabling identification of duplex groups (DGs), non-overlapping groups (NGs), and triplex groups (TGs). The final data is available in CRIS for downstream exploration, further complimented by visualization tools such as IGV (read-level inspection), VARNA (2D structure rendering, PyMOL (3D modeling), and crosslink-focused utilities like *nt_freq.py* and *freq_icSHAPE.py*. This allows for a thorough analysis of RNA structure, dynamics, and phylogeny, as well as interactions derived from contact data.

To reduce the storage footprint of XLRNA sequencing data, we developed bam2bedz, a lossy compression tool tailored to extract essential information from RNA crosslinking alignments. This approach selectively extracts the read start position, CIGAR string, MD tag, read count, and strand information for simple gap-typed reads (prigap1, prigapm, prihomo) and stores the data in a compressed, modified BED format. For trans-gapped reads (pritrans), the two RNA arms are recorded in a modified BEDPE file that retains the chromosome and start position of each arm, CIGAR string, MD tag, read count, and strand information for both arms. Continuous reads (pricont), which resemble conventional RNA-seq coverage profiles, are processed and stored as compressed bigWig files that track per-base coverage. By selectively retaining only the most relevant features, bam2bedz significantly reduces data size while maintaining analytical integrity.

Unlike reference-based formats like CRAM, bam2bedz outputs eliminates redundancy from full sequences and removes reference reconstruction for visualization (60). Since the XLRNA approach for structural visualization relies on high fidelity string matches rather than per-base sequence information, bam2bedz retains only positional and topological information. The resulting “bed.gz” and “bedpe.gz” compressed files are lightweight, tab-delimited, and human-readable, allowing rapid inspection and manipulation using tools such as bedtools, awk, pandas, and tabix (for fast querying via indexes). Critically, the compressed files can readily be expanded if needed, using the original reference genome to restore alignment-level information for most structural analyses, providing flexibility for both storage and interpretation.

To empirically benchmark compression efficiency, we applied bam2bedz to multiple datasets (e.g., PARIS, SHARC), sampled increasing numbers of mapped reads (e.g., 1 million to 100 million) using samtools and shuf, then compressed them using bam2bedz. For all sampled data, bam2bedz achieved consistent and substantial reductions in file size across all XLRNA reads (**Supplemental Figure 1**). Compression efficiency was assessed by calculating the bytes per read (BPR) for both BAM and BED outputs, excluding BAM headers to ensure a direct comparison between the original and compressed output.

Since pricont, prigap1, and pritrans BAM files are the largest files in a typical XLRNA dataset, we benchmarked bam2bedz on subsets of these data. For prigap1, 1000 randomized rads from PARIS data containing read name with experimental conditions, tags, CIGAR strings, and alignment metadata, average ∼350 BPR. The equivalent BED lines contain ∼30 BPR, resulting in an approximate 12-fold reduction in BPR where reads are unique and contain few or no overlaps. Similarly, other datasets in CRIS show the expected ∼12-fold reduction, since reads are short and are expected to have mildly repetitive regions. For synthetic datasets with repetitive sequences or where sequencing coverage is near saturation such that every position in the genome is overlapped by multiple reads, we observe up to ∼20-fold reduction in size.

### Genome References

CRIS provides three categories of reference genomes that support accurate and consistent alignment of sequencing data across research applications.

1. “Standard”: Unmodified genomes for human (hg38) and mouse (mm10) are linked, ensuring compatibility with common RNA research pipelines.
2. “Curated”: Repetitive-element masked genomes to improve mapping accuracy (37). Effectively a modified reference for studying noncoding and repetitive RNA sequences.
3. “Special:” Custom designed genomes for RNA-RNA interaction studies. Manual preparation of concatenated interacting RNA genes into a single reference file, allowing mapping analysis of complex RNA-RNA interactions within structured multi-gene contexts.

All references are fully downloadable via CRIS with companion STAR alignment indices.

### Data Quality Control

The currently published, pre-processed datasets were first subjected to FastQC analysis to assess sequencing quality, overrepresented sequences, and adapter content. The resulting FastQC reports are accessible via the “Data” page, which also includes plots of gap and segment length distributions, hallmark measures of crosslinking data quality. A comprehensive statistics report was also generated, extracting and aggregating sequencing and alignment metrics from the processed data. Read counts from raw FASTQ files, alignment metrics from the Log.final.out files generated by STAR, and gap analysis results from SLURM output logs were consolidated into a structured CSV file, ensuring consistent formatting for downstream analysis and reporting. The sequencing statistics report enables reproducible, scalable assessment of sequencing quality, mapping performance, and gap-type distributions.

As part of the initiative to standardize the processing of XLRNA data in CRSSANT analysis, CRIS integrates “genes2bed,” a script designed for filtering genes used in DG clustering by read count thresholds and extracting their corresponding genomic intervals in BED format. The data processing pipeline guarantees that structural and interactional insights are supported by robust experimental evidence, since the DGs are formed based on existing read information. This tool ultimately simplifies the process of clustering RNA duplexes and enables users to process data without needing to manually prepare the BED file necessary for CRSSANT.

### Data Visualization

Integration of XLRNA data with genome browsers like the Integrative Genomics Viewer (IGV) is not streamlined, lacks clear guidelines for data visualization, local feature extraction, and overlay with conservation tracks (61). Introduced as part of the CRIS toolkit, the updated bedpe2arc12 script enhances the visualization of DGs identified by CRSSANT by converting bedpe records into bed12 format for arc-based display. Unlike its predecessor scripts dg2arc.py and bedpetobed12.py, this version resolves alignment conversion issues and introduces a dynamic RGB coloring scheme to reflect DG score intensity (**Figure 2a, b**). Specifically, the script processes each DG independently, cycling through a predefined set of base colors to assign a unique hue per group of DGs. Each group of DGs are sorted by raw score, and their color intensity is modulated using a gamma-corrected log transformation of the score (**Figure 2c, d**). This results in higher-scoring DGs appearing darker, drawing visual attention to significant interactions. In addition to coloring, the script ensures strand concordance and precise block coordinate mapping, enabling accurate spatial representation of RNA-RNA interactions. Altogether, these improvements support more intuitive data interpretation and facilitate the prioritization of high-confidence DGs in large-scale structural studies. The script requires only an input bedpe file that summarizes CRSSANT DGs and an output name for the bed12 file containing the arcs.

**Figure 2.**
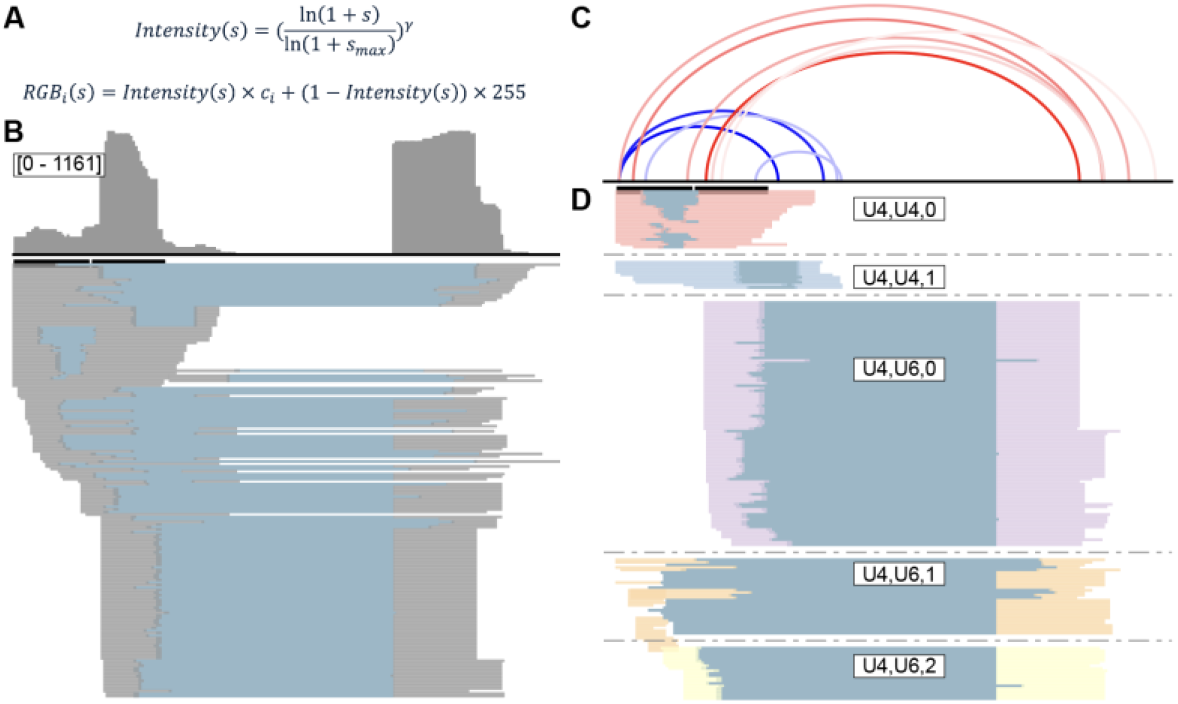
Visualization of RNA duplex groups (DGs) using the bedpe2arc12 script from the CRIS toolkit. (A) DG color intensity is computed using a gamma-corrected log transformation of each RGB color channel c_i_, where s is the raw DG score for the pair, s_max_ is the highest score in the group, and γ is the gamma correction factor (default: 0.5). (B) Standard genome browser view showing per-base coverage (top) and read pair alignments (gray/blue). While informative, this raw format lacks explicit interaction grouping and becomes visually cluttered in high-coverage regions. (C) Enhanced arc rendering using bedpe2arc12, showing score-weighted arcs and group-wise coloring (e.g., U4-U4 in blue, U4-U6 in red). Higher-scored, confident DGs appear darker. (D) Block-level alignments for DG pairs, each corresponding to a distinct DG subgroup, supporting comparison across alternative structures or interaction variants.

### CRIS Enables High-resolution, Protocol-agnostic RNA Analysis

A central challenge in transcriptome-wide RNA structure analysis lies in integrating data from distinct methodologies, each with its own chemistries, biological systems, biases, resolution, and structural assumptions. CRIS addresses this challenge by passing crosslinking data through a standardized computational pipeline, enabling rigorous comparison across methods. Moreover, the approach enables chemical probing data to be neatly integrated.

### Transcriptome-wide DG Analysis

Using data from PARIS studies, CRIS reveals thousands of RNA duplexes across the entire transcriptome. Here, we highlight extracted data on XIST, a long noncoding RNA known for its complex regulatory roles and architectural modularity, as a model transcript for cross-platform structural analysis (Figure 4). CRIS integrates PARIS (long-range interactions) and icSHAPE (local flexibility) data into a single visualization step, resolving a longstanding comparison obstacle.

SHAPE-based methods yield nucleotide-resolution reactivity scores that indirectly reflect RNA secondary structure, but they neither resolve long-range or noncanonical interactions, nor directly report on duplex

formation. In contrast, XLRNA techniques like PARIS physically capture base-pairing interactions, generating duplex maps that span large sequence distances. By mapping both data types to the same transcript and presenting them as aligned genome browser tracks, CRIS removes format disparities and enables accurate positional overlay and comparison (**Figure 3A**). This eliminates ambiguity in interpreting structural signals and enables joint solving of local and global RNA architectures. Using CRSSANT analysis, CRIS reveals modular and highly structured intra-exonic duplexes, many coinciding with drops in icSHAPE reactivity (**Figure 3B**). This is consistent with base-pairing and structural domains identified by CRIS (**Figure 3C**), which delineate folding assumptions and reflect the hierarchical organization of XIST required X-chromosome silencing. Duplexes cleanly align with peaks in PARIS read coverage (**Figure 3D**), further supporting their structural relevance.

**Figure 3.**
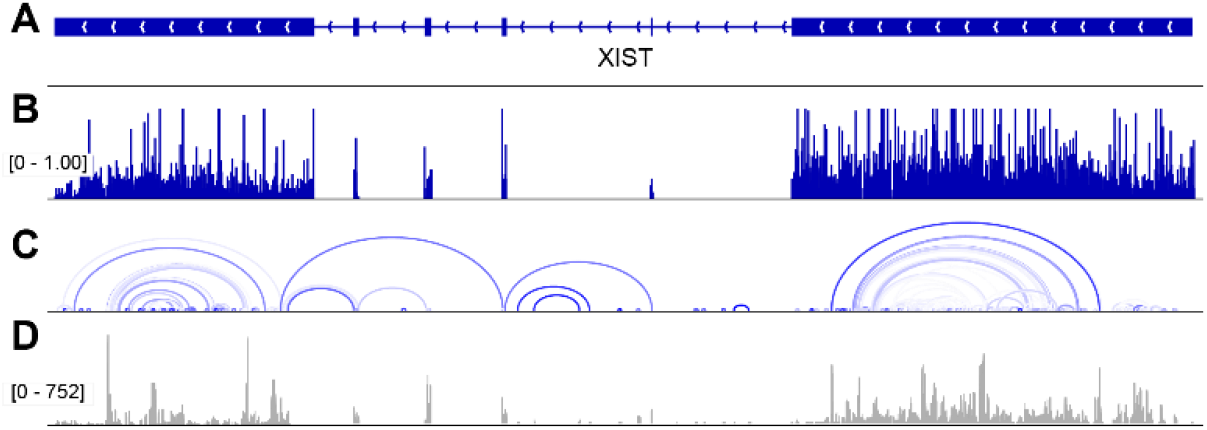
Multitrack genome browser-style track alignment facilitates cross-comparison of complementary representation of RNA structural features. (A) Exonic regions of the XIST locus as thick lines. (B) icSHAPE reactivity profile plotted as nucleotide-level scores (0 - 1.0) indicating single-strandedness and structural flexibility. (C) Duplex group arcs identified by CRSSANT, representing captured RNA-RNA base-pairing interactions along the XIST transcript; the span and density of arcs reflect the complexity of molecular interactions complemented by SHAPE data. Higher-scored, confident DGs appear darker. (D) PARIS sequencing coverage showing read depth (0 - 752 reads), highlighting regions with high RNA-RNA contact frequency.

While icSHAPE data alone may suggest regions of low flexibility, only XLRNA data can determine whether these correspond to local helices or distal interactions that form tertiary structures, which is a critical distinction for understanding RNA function. More broadly, this analysis exemplifies CRIS as a scalable, multi-modal framework for structural inference using both chemical probing and crosslinking data. Previous approaches relied on qualitative comparisons, overlaying reactivity scores, and structural predictions without grounding them in empirically verified long-range interactions. CRIS enables identification of structurally insulated regions, cooperative domains, and potential tertiary motifs with minimal effort.

### Comparative Analysis of Protocols

Another example of CRIS-processed data used to contrast methodologies is the direct comparison of RNA duplexes identified by PARIS1 and STAU1 hiCLIP in the SRSF1 exon 4 region (**Figure 4**). These datasets represent fundamentally different strategies for probing RNA structure: PARIS captures transcriptome-wide RNA duplexes without protein bias, while hiCLIP targets RNA structures bound specifically by STAU1, a double-stranded RNA-binding protein (dsRBP).

**Figure 4.**
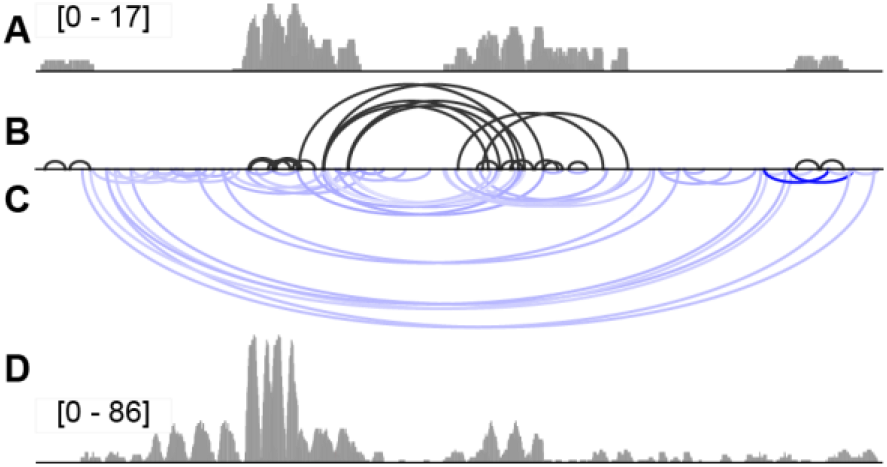
Comparison of RNA duplexes identified by PARIS and STAU1 hiCLIP data in the SRSF1 exon 4 region. (A) Genomic alignment coverage of STAU1 hiCLIP duplex groups, highlighting specific dsRBP-bound RNA structures. (B) STAU1 hiCLIP duplex groups, with data combined from three independent sequencing libraries and visualized as arcs. (C) Arc diagram of PARIS1 duplex groups, highlighting long-range interactions and structural complexity, vertically inverted for direct comparison (D) Coverage of RNA duplex groups captured using PARIS1 in HeLa cells, aligned to the transcriptome. PARIS1 captures all previously known STAU1-bound duplexes and reveals additional RNA structures undetected by hiCLIP.

Comparing PARIS and hiCLIP datasets is inherently challenging due to differences in experimental design, resolution, and mapping assumptions. However, processing both through the CRIS pipeline normalizes variables, allowing for standardization of data structure, clustering parameters, and genomic alignment, all to enable a fair comparison. CRIS ensures that duplexes from both PARIS and hiCLIP are grouped, quantified, and visualized within the same computational framework, minimizing technical variability and highlighting true biological differences. While CLIP-based methods provide insight into individual protein binding, they do not capture a global view of RNA structure or binding context (**Figure 4A**). In contrast, PARIS-like approaches detect RNA duplexes transcriptome-wide, enabling protein-agonistic structural analysis and provide a comprehensive view of the RNA (**Figure 4D**). Here, we show that PARIS not only recapitulates all duplexes detected by STAU1 hiCLIP but also identifies a broader set of additional duplexes in the same region (**Figure 4B, 4C**). These additional interactions likely represent other dsRBP targets or functional RNA structures that escape CLIP detection due to transient binding or low affinity. This complementary and comprehensive coverage demonstrates how CRIS enables cross-platform integration, providing a richer and more complete view RNA structure landscapes.

Unifying PARIS and CLIP data at the RNA duplex group level represents a major improvement to the analysis workflow. Historically, comparisons between structural and CLIP-based data were qualitative, often limited to overlaying raw read densities or peak annotations. CRIS enables quantitative, interpretable structural comparisons at the duplex level, showing not just whether a region is structured, but how structures align or diverge across methods. This also reveals broader structural context around dsRBP binding and supports hypothesis generation about cooperative or competitive binding by other factors. For instance, the additional duplexes detected only by PARIS suggest possible alternative binding sites, structural switches, or regulatory elements obscured in the protein-centric view provided by CLIP.

### Discovery of Alternative Conformations

Small nucleolar RNAs (snoRNAs) often adopt complex secondary and tertiary structures essential for their function in ribosomal RNA (rRNA) maturation (62–64). Data from PARIS2 experiments demonstrated that U8 snoRNA, previously thought to adopt a single structure, exists in multiple conformational states, including a “linear precursor” form and a “maturely folded” structure required for its biological activity in the nucleolus. CRIS data resolves multiple alternative conformations of the U8 snoRNA. DG clustering combined with 3D distance constraints reveals the presence of conformational subpopulations, suggesting dynamic structural regulation (**Figure 5**). This confirms that U8 adopts distinct conformations, consistent with the hypothesis of structure-dependent processing. More importantly, they demonstrate how CRIS enables unsupervised structural inference at the group level, rather than relying on predefined RNA folding rules or annotations. Similar approaches were applied to other snoRNAs and viral RNAs (34,35,62).

**Figure 5.**
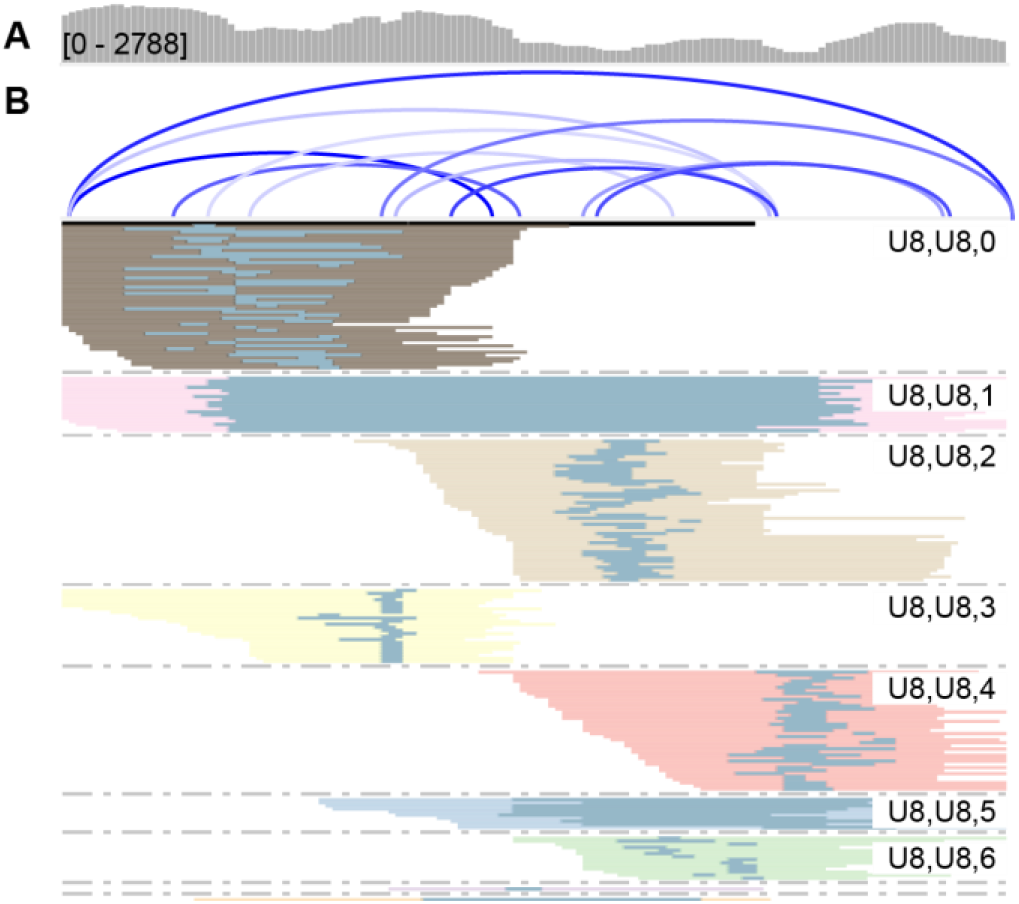
Example of U8 analysis. (A) Arc plots show distinct sets of duplex interactions, separated into groups using CRSSANT clustering. (B) DGS support an extended, linear structure consistent with the transient precursor form. (C) Dimer base pairs or intermolecular overlapping DGs recapitulate the “closed-loop” base-pairing pattern observed in mature U8. (D) Secondary structure rendering depicts spatially distinct conformations supported by CRIS-inferred base-pairing interactions. This example highlights the capacity of CRIS to resolve structural heterogeneity within a single RNA species, revealing dynamic conformational states that are otherwise difficult to distinguish using traditional secondary structure prediction or ensemble models alone. By leveraging crosslink-informed clustering, CRIS allows researchers to explore alternative structural landscapes, quantify supporting sequencing data, and visualize them in a way that integrates naturally with 3D modeling tools.

## METHODS

Data pre-processing, read alignment, and statistical analysis were performed following the CRSSANT pipeline (37,59). Initially, all raw FASTQ files were inspected with FastQC to assess sequencing quality and adapter content (65). Where appropriate, adapter sequences were trimmed from the 3’ ends using Trimmomatic, and multiplexed libraries were demultiplexed and split by barcode using splitFastq.pl from the icSHAPE pipeline (30,49,66). PCR duplicates were removed using readCollapse, after which a second round of Trimmomatic trimming was completed to remove residual 5’ adapter sequences.

Reads were aligned using STAR v2.7.1a with permissive alignment parameters to maximize sensitivity for chimeric reads (67). Primary alignments were extracted and initially classified into gap types using gaptypes.py from the CRSSANT pipeline (37). Reads identified as continuous were further examined for soft-clipped regions and potential reverse chimeras. These were processed with softreverse.py and remapped using STAR, followed by a second extraction and classification of primary alignments. Outputs from both rounds of mapping were merged and reads with splice junctions or short deletions (1-2 nt gaps) were filtered out using gapfilter.py to refine the prigap1 and prigapm datasets.

Finally, updated gaplendist.py and seglendist.py scripts from the rna2d3d repository were used to plot the gap-length and segment-length distributions, respectively (59). These distributions, along with summary statistics, were recorded for each dataset using a wrapper script to evaluate crosslinking quality and structural features of RNA interactions.

## CONCLUSIONS

The Crosslinking-based RNA Interactomes and Structurome (CRIS) database makes a substantial contribution to the RNA research community by offering rigorously analyzed, pre-processed RNA structure and interaction datasets. Prioritizing data quality, transparency, and usability, CRIS addresses challenges like inconsistent quality control and the computational demands typically associated with raw data analysis. Its intuitive interface, detailed documentation, and ready-to-use datasets allow researchers to focus directly on biological interpretation and hypothesis generation without requiring prior coding knowledge or heavy computational resources.

By unifying disparate pipelines and XLRNA data, CRIS enables previously impractical quantitative and qualitative comparisons, accelerating discoveries in RNA-based drug design and synthetic biology. Its emphasis on transparency and data integrity highlights its reliability and promotes broader accessibility within the research community.

At the same time, it further serves as a training platform for machine learning models. CRIS has the potential to support both supervised and unsupervised learning tasks. This platform supports a range of supervised learning tasks, including classification of conformations and interaction types, as well as regression-based methods in predicting structural stability and interaction strength. In unsupervised learning settings, CRIS enables clustering of RNA interaction hotspots based on shared structural or sequence-based features, and can facilitate dimensionality reduction techniques (e.g., PCA, t-SNE, UMAP) for visualization and embedding of high-dimensional RNA data (68,69). Additionally, the diversity of cell types, species, and experimental conditions captured in the database makes it ideal for transfer learning and multi-task learning approaches already implemented in other sequencing techniques, helping to build models with broader applicability to biological systems (70).

CRIS also bridges a wide range of artificial intelligence tasks. For instance, its datasets can be used to train deep learning models that predict RNA secondary and tertiary structures directly from sequence data, infer complex RNA-RNA interaction networks, or identify disease-relevant structures (43,71,72). CRIS can also enable unsupervised AI tasks, such as embedding structural features into latent spaces or learning representations of RNA characteristics using autoencoders or contrastive learning (73). Future applications include support for graph neural networks for learning from RNA structure graphs, as well as transformer-based models for sequence-to-structure prediction (44).

Designed as a dynamic and evolving platform, CRIS continually expands its features, datasets, and analytical capabilities to meet emerging research needs. Planned enhancements aim to improve user experience and data exploration through tools like interactive tables summarizing key RNA-RNA interactions, additional data collection to broaden species and cell-type coverage. While the current data highlights crosslink-ligation RNA sequencing datasets, future updates will integrate data such as protein-RNA interactions to support complex biological inquiries, and we hope that establishing the CRIS pipeline will support cross-study comparisons. Future iterations of the CRIS database will incorporate information on RNA splicing, editing, and other processing events, offering a more comprehensive view of RNA biology. Continuous refinement of data curation standards and computational tools ensures the platform remains a relevant and valuable resource.

As RNA research evolves, addressing challenges like multi-omics integration, dynamic RNA structure analysis, and large-scale data handling will require adaptable tools. CRIS is committed to meeting these demands, serving as both a foundational database and a catalyst for future discoveries in RNA biology and therapeutics.

## DATA AVAILABILITY

CRIS is accessible at https://whl-usc.github.io/cris/home. Published sequencing data can be found on NCBI GEO and the ENA Browser, while processed data are available for download via the links on the “Data” page of the CRIS website.

## ACKNOWLEDGEMENTS

We thank current members of the Lu Lab for their assistance with data processing and software testing, and former members for their preparation of experimental data. We are especially grateful to intern Casper Peters and volunteer Jennifer Galvez for their feedback on the CRIS website design and user experience. We also thank members of the USC Alfred E. Mann School of Pharmacy and Pharmaceutical Sciences, including collaborators Dr. Mario Alba and Brittney Hua, for their valuable feedback during the testing of the CRIS database.

## CONTRIBUTIONS

W.H.L. and Z.L. conceived and designed the project. Z.L., M.Z., and W.H.L. developed the methodology. Software development and deployment was carried out by W.H.L., P.S., and M.Z. Data generation was performed by K.L., J.B., M.Z., and W.H.L. W.H.L., A.S., and C.D. curated and organized the data. Validation was overseen by W.H.L. and Z.L. W.H.L. and C.D. wrote the manuscript. All authors contributed to reviewing and editing. Data visualization was performed by W.H.L. and C.D. Project supervision was provided by Z.L. and W.H.L., and funding was acquired by Z.L.

## SUPPLEMENTARY DATA

**Supplemental Figure 1.**
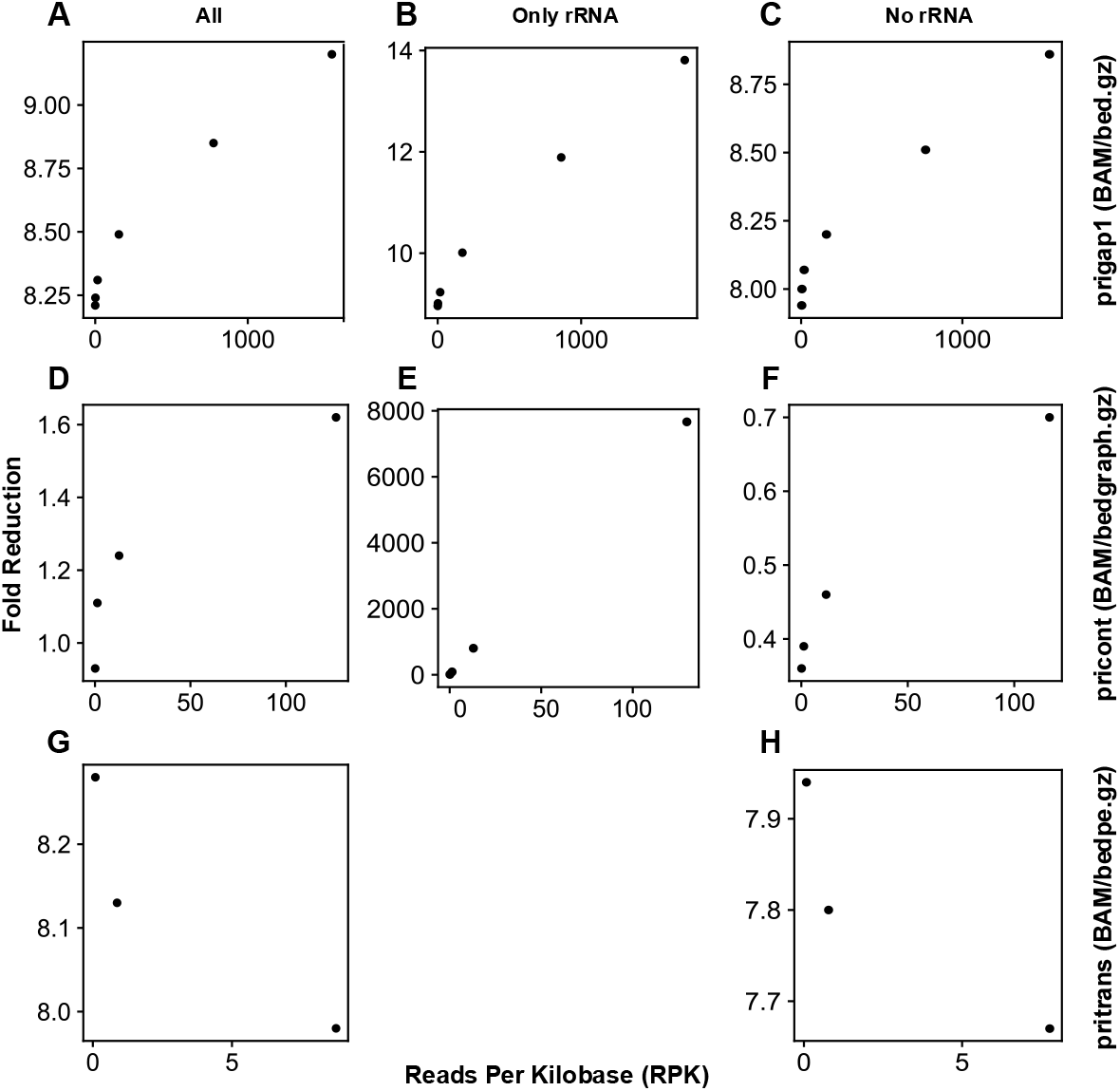
Compression performance of bam2bedz across read types and genomic content. Plots show the relationship between RPK (Reads Per Kilobase) and file size reduction for different classes of RNA-seq reads after bam2bedz compression. Each subpanel depicts trends across increasing read numbers for: (A) all prigap1 reads, (B) prigap1 reads mapping exclusively to rRNA regions, and (C) prigap1 reads excluding rRNA. Compression scales logarithmically with RPK, reflecting increasing alignment redundancy. (E, F) pricont reads show limited compression (∼3-fold BPR) due to retention of position specific details in BEDgraph format. (E) pricont rRNA-only reads, by contrast, compress more efficiently and linearly with read count because all alignments localize to a single pseudo-chromosome in the STAR-masked genome. (G, H) pritrans reads represent dual-arm, long-range interactions and require structural pairing during compression. This category shows a log-decay compression profile, achieving up to ∼8-fold BPR with ∼1 million reads.

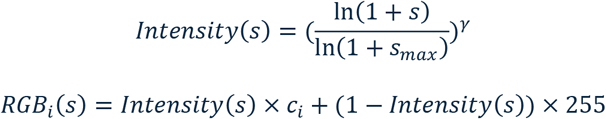

Measurement of bam2bedz efficiency can be visualized by plotting RPK (reads per kilobase) against compression ratio, where:

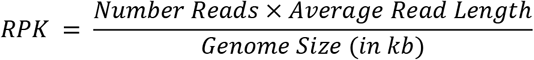

High RPK values indicate denser coverage, which correlates with greater compression due to repeated alignments. In the example of SHARC data compression, 10 million ∼50-nt reads mapped to the hg38 reference genome (∼3.1 million kb) yields an RPK of approximately 155, suggesting low genomic saturation and high compressibility Decreasing the number of reads dramatically increases the uniqueness of each read, therefore reducing the compression ratio. The same logic can be applied to prigapm and prihomo reads, showing a decrease in file size that plateaus for higher read numbers. When processing only ribosomal RNA (rRNA) reads, we note an increase in compression, up to ∼13-fold BPR, due to the nature of repeated alignments in the densely covered rRNA regions.

In contrast, non-rRNA reads scale less favorably, with compression not exceeding 9-fold for up to 100 million reads (∼1500 RPK). Compression scales logarithmically with read depth due to increased alignment redundancy but plateaus as data entropy declines and redundancy is exhausted. Compression of pricont reads follows a similar trend of logarithmic increase in compression, not exceeding ∼3-fold BPR for combined reads. This limited gain is expected, as reads mapping to distinct chromosomes retain separate coverage tracks in the BEDgraph format, preserving chromosome-specific detail. In contrast, rRNA-only reads compress more efficiently and scale linearly with read count, since all alignments are confined to a single “chromosome” in the custom STAR-masked genome used for mapping.

Pritrans reads, which capture long-range or inter-molecular RNA interactions, require a distinct compression strategy due to their dual-arm structure. Unlike simple gapped or continuous reads that map to a single genomic locus, pritrans reads involve two separate alignments, each corresponding to one RNA arm on different chromosomes or distant loci within the same transcript. By compressing both arms into a single structured record, bam2bedz retains topological integrity essential for downstream structure inference, clustering, and visualization of long-range RNA contacts. The data exhibits a log decay pattern in file size compression, with ∼1 million reads reducing ∼8-fold BPR.

## REEFERENCES

1. Das R. RNA structure: a renaissance begins? Nat Methods. 2021 May;18(5):439–439. doi:10.1038/s41592-021-01132-4

2. Gebert LFR, MacRae IJ. Regulation of microRNA function in animals. Nat Rev Mol Cell Biol. 2019 Jan;20(1):21–37. doi:10.1038/s41580-018-0045-7 PubMed PMID: 30108335; PubMed Central PMCID: PMC6546304.

3. Bratkovič T, Božič J, Rogelj B. Functional diversity of small nucleolar RNAs. Nucleic Acids Res. 2020 Feb 28;48(4):1627–51. doi:10.1093/nar/gkz1140

4. Bridges MC, Daulagala AC, Kourtidis A. LNCcation: lncRNA localization and function. J Cell Biol. 2021 Feb 1;220(2):e202009045. doi:10.1083/jcb.202009045 PubMed PMID: 33464299; PubMed Central PMCID: PMC7816648.

5. Mortimer SA, Kidwell MA, Doudna JA. Insights into RNA structure and function from genome-wide studies. Nat Rev Genet. 2014 Jul;15(7):469–79. doi:10.1038/nrg3681

6. Loda A, Heard E. Xist RNA in action: Past, present, and future. PLoS Genet. 2019 Sep;15(9):e1008333. doi:10.1371/journal.pgen.1008333 PubMed PMID: 31537017; PubMed Central PMCID: PMC6752956.

7. Kudla G, Granneman S, Hahn D, Beggs JD, Tollervey D. Cross-linking, ligation, and sequencing of hybrids reveals RNA–RNA interactions in yeast. Proc Natl Acad Sci. 2011 Jun 14;108(24):10010–5. doi:10.1073/pnas.1017386108

8. Juzwik CA, S Drake S, Zhang Y, Paradis-Isler N, Sylvester A, Amar-Zifkin A, et al. microRNA dysregulation in neurodegenerative diseases: A systematic review. Prog Neurobiol. 2019 Nov;182:101664. doi:10.1016/j.pneurobio.2019.101664 PubMed PMID: 31356849.

9. Wang XW, Liu CX, Chen LL, Zhang QC. RNA structure probing uncovers RNA structure-dependent biological functions. Nat Chem Biol. 2021 Jul;17(7):755–66. doi:10.1038/s41589-021-00805-7

10. Zhou LY, Qin Z, Zhu YH, He ZY, Xu T. Current RNA-based Therapeutics in Clinical Trials. Curr Gene Ther. 19(3):172–96. doi:10.2174/1566523219666190719100526

11. Zhu Y, Zhu L, Wang X, Jin H. RNA-based therapeutics: an overview and prospectus. Cell Death Dis. 2022 Jul 23;13(7):1–15. doi:10.1038/s41419-022-05075-2

12. Chen J, Lin J, Hu Y, Ye M, Yao L, Wu L, et al. RNADisease v4.0: an updated resource of RNA-associated diseases, providing RNA-disease analysis, enrichment and prediction. Nucleic Acids Res. 2023 Jan 6;51(D1):D1397–404. doi:10.1093/nar/gkac814 PubMed PMID: 36134718; PubMed Central PMCID: PMC9825423.

13. Thomas SJ, Moreira ED, Kitchin N, Absalon J, Gurtman A, Lockhart S, et al. Safety and Efficacy of the BNT162b2 mRNA Covid-19 Vaccine through 6 Months. N Engl J Med. 2021 Nov 3;385(19):1761– 73. doi:10.1056/NEJMoa2110345

14. Sahin U, Muik A, Vogler I, Derhovanessian E, Kranz LM, Vormehr M, et al. BNT162b2 vaccine induces neutralizing antibodies and poly-specific T cells in humans. Nature. 2021 Jul;595(7868):572– 7. doi:10.1038/s41586-021-03653-6

15. Wang S, Li H, Lian Z, Deng S. The Role of RNA Modification in HIV-1 Infection. Int J Mol Sci. 2022 Jul 8;23(14):7571. doi:10.3390/ijms23147571 PubMed PMID: 35886919; PubMed Central PMCID: PMC9317671.

16. Yu H, Wen Y, Yu W, Lu L, Yang Y, Liu C, et al. Optimized circular RNA vaccines for superior cancer immunotherapy. Theranostics. 2025;15(4):1420–38. doi:10.7150/thno.104698 PubMed PMID: 39816687; PubMed Central PMCID: PMC11729565.

17. Ratni H, Ebeling M, Baird J, Bendels S, Bylund J, Chen KS, et al. Discovery of Risdiplam, a Selective Survival of Motor Neuron-2 (SMN2) Gene Splicing Modifier for the Treatment of Spinal Muscular Atrophy (SMA). J Med Chem. 2018 Aug 9;61(15):6501–17. doi:10.1021/acs.jmedchem.8b00741

18. Piñeiro D, Martinez-Salas E. RNA Structural Elements of Hepatitis C Virus Controlling Viral RNA Translation and the Implications for Viral Pathogenesis. Viruses. 2012 Oct 19;4(10):2233–50. doi:10.3390/v4102233 PubMed PMID: 23202462; PubMed Central PMCID: PMC3497050.

19. Burrill CP, Westesson O, Schulte MB, Strings VR, Segal M, Andino R. Global RNA Structure Analysis of Poliovirus Identifies a Conserved RNA Structure Involved in Viral Replication and Infectivity. J Virol. 2013 Nov;87(21):11670–83. doi:10.1128/JVI.01560-13 PubMed PMID: 23966409; PubMed Central PMCID: PMC3807356.

20. Zhu Y, Chaubey B, Olsen GL, Varani G. Structure of Essential RNA Regulatory Elements in the West Nile Virus 3′-Terminal Stem Loop. J Mol Biol. 2024 Nov;436(22):168767. doi:10.1016/j.jmb.2024.168767

21. Gote V, Bolla PK, Kommineni N, Butreddy A, Nukala PK, Palakurthi SS, et al. A Comprehensive Review of mRNA Vaccines. Int J Mol Sci. 2023 Jan;24(3):2700. doi:10.3390/ijms24032700

22. Lu Z, Chang HY. The RNA Base-Pairing Problem and Base-Pairing Solutions. Cold Spring Harb Perspect Biol. 2018 Dec 1;10(12):a034926. doi:10.1101/cshperspect.a034926 PubMed PMID: 30510063.

23. Schroeder SJ. Challenges and approaches to predicting RNA with multiple functional structures. RNA N Y N. 2018 Dec;24(12):1615–24. doi:10.1261/rna.067827.118 PubMed PMID: 30143552; PubMed Central PMCID: PMC6239171.

24. Bugnon LA, Edera AA, Prochetto S, Gerard M, Raad J, Fenoy E, et al. Secondary structure prediction of long noncoding RNA: review and experimental comparison of existing approaches. Brief Bioinform. 2022 Jul 18;23(4):bbac205. doi:10.1093/bib/bbac205 PubMed PMID: 35692094.

25. Ma H, Jia X, Zhang K, Su Z. Cryo-EM advances in RNA structure determination. Signal Transduct Target Ther. 2022 Feb 23;7(1):1–6. doi:10.1038/s41392-022-00916-0

26. Jackson RW, Smathers CM, Robart AR. General Strategies for RNA X-ray Crystallography. Mol Basel Switz. 2023 Feb 23;28(5):2111. doi:10.3390/molecules28052111 PubMed PMID: 36903357; PubMed Central PMCID: PMC10004510.

27. Velema WA, Lu Z. Chemical RNA Cross-Linking: Mechanisms, Computational Analysis, and Biological Applications. JACS Au. 2023 Feb 27;3(2):316–32. doi:10.1021/jacsau.2c00625

28. Marušič M, Toplishek M, Plavec J. NMR of RNA - Structure and interactions. Curr Opin Struct Biol. 2023 Apr 1;79:102532. doi:10.1016/j.sbi.2023.102532

29. Kotar A, Foley HN, Baughman KM, Keane SC. Advanced approaches for elucidating structures of large RNAs using NMR spectroscopy and complementary methods. Methods. 2020 Nov 1;Methods to characterize virus small RNAs and RNA structures 183:93–107. doi:10.1016/j.ymeth.2020.01.009

30. Flynn RA, Zhang QC, Spitale RC, Lee B, Mumbach MR, Chang HY. Transcriptome-wide interrogation of RNA secondary structure in living cells with icSHAPE. Nat Protoc. 2016 Feb;11(2):273–90. doi:10.1038/nprot.2016.011

31. Merino EJ, Wilkinson KA, Coughlan JL, Weeks KM. RNA Structure Analysis at Single Nucleotide Resolution by Selective 2’-Hydroxyl Acylation and Primer Extension (SHAPE). J Am Chem Soc. 2005 Mar 1;127(12):4223–31. doi:10.1021/ja043822v

32. Spitale RC, Crisalli P, Flynn RA, Torre EA, Kool ET, Chang HY. RNA SHAPE analysis in living cells. Nat Chem Biol. 2013 Jan;9(1):18–20. doi:10.1038/nchembio.1131 PubMed PMID: 23178934; PubMed Central PMCID: PMC3706714.

33. Wilkinson KA, Merino EJ, Weeks KM. Selective 2′-hydroxyl acylation analyzed by primer extension (SHAPE): quantitative RNA structure analysis at single nucleotide resolution. Nat Protoc. 2006 Aug;1(3):1610–6. doi:10.1038/nprot.2006.249

34. Zhang M, Li K, Bai J, Velema WA, Yu C, van Damme R, et al. Optimized photochemistry enables efficient analysis of dynamic RNA structuromes and interactomes in genetic and infectious diseases. Nat Commun. 2021 Apr 20;12(1):2344. doi:10.1038/s41467-021-22552-y

35. Van Damme R, Li K, Zhang M, Bai J, Lee WH, Yesselman JD, et al. Chemical reversible crosslinking enables measurement of RNA 3D distances and alternative conformations in cells. Nat Commun. 2022 Feb 17;13(1):911. doi:10.1038/s41467-022-28602-3

36. Lee WH, Li K, Lu Z. Chemical crosslinking and ligation methods for in vivo analysis of RNA structures and interactions. In: Yamagami R, Kwok CK, editors. Methods in Enzymology [Internet]. Academic Press; 2023 [cited 2025 Feb 12]. p. 253–81. (Enzymes in RNA Science and Biotechnology Part A). Available from: https://www.sciencedirect.com/science/article/pii/S0076687923000873 doi:10.1016/bs.mie.2023.02.020

37. Zhang M, Hwang IT, Li K, Bai J, Chen JF, Weissman T, et al. Classification and clustering of RNA crosslink-ligation data reveal complex structures and homodimers. Genome Res. 2022 May;32(5):968–85. doi:10.1101/gr.275979.121 PubMed PMID: 35332099; PubMed Central PMCID: PMC9104705.

38. Zhou J, Li P, Zeng W, Ma W, Lu Z, Jiang R, et al. IRIS: A method for predicting in vivo RNA secondary structures using PARIS data. Quant Biol. 2020;8(4):369–81. doi:10.1007/s40484-020-0223-4

39. Kozar I, Philippidou D, Margue C, Gay LA, Renne R, Kreis S. Cross-Linking Ligation and Sequencing of Hybrids (qCLASH) Reveals an Unpredicted miRNA Targetome in Melanoma Cells. Cancers. 2021 Mar 4;13(5):1096. doi:10.3390/cancers13051096 PubMed PMID: 33806450; PubMed Central PMCID: PMC7961530.

40. Andrews RJ, Baber L, Moss WN. RNAStructuromeDB: A genome-wide database for RNA structural inference. Sci Rep. 2017 Dec 8;7(1):17269. doi:10.1038/s41598-017-17510-y

41. Zuker M. Mfold web server for nucleic acid folding and hybridization prediction. Nucleic Acids Res. 2003 Jul 1;31(13):3406–15. doi:10.1093/nar/gkg595

42. Reuter JS, Mathews DH. RNAstructure: software for RNA secondary structure prediction and analysis. BMC Bioinformatics. 2010 Mar 15;11(1):129. doi:10.1186/1471-2105-11-129

43. Townshend RJL, Eismann S, Watkins AM, Rangan R, Karelina M, Das R, et al. Geometric deep learning of RNA structure. Science. 2021 Aug 27;373(6558):1047–51. doi:10.1126/science.abe5650

44. Shen T, Hu Z, Sun S, Liu D, Wong F, Wang J, et al. Accurate RNA 3D structure prediction using a language model-based deep learning approach. Nat Methods. 2024 Dec;21(12):2287–98. doi:10.1038/s41592-024-02487-0

45. Gong J, Shao D, Xu K, Lu Z, Lu ZJ, Yang YT, et al. RISE: a database of RNA interactome from sequencing experiments. Nucleic Acids Res. 2018 Jan 4;46(Database issue):D194–201. doi:10.1093/nar/gkx864 PubMed PMID: 29040625; PubMed Central PMCID: PMC5753368.

46. Charting targeted courses for vaccination. Nat Biomed Eng. 2025 Feb;9(2):149–50. doi:10.1038/s41551-025-01366-z

47. Mahmoudian F, Ahmari A, Shabani S, Sadeghi B, Fahimirad S, Fattahi F. Aptamers as an approach to targeted cancer therapy. Cancer Cell Int. 2024 Mar 16;24(1):108. doi:10.1186/s12935-024-03295-4

48. Mullard A. FDA approves second RNA aptamer. Nat Rev Drug Discov. 2023 Sep 6;22(10):774–774. doi:10.1038/d41573-023-00148-z

49. Lu Z, Zhang QC, Lee B, Flynn RA, Smith MA, Robinson JT, et al. RNA Duplex Map in Living Cells Reveals Higher-Order Transcriptome Structure. Cell. 2016 May 19;165(5):1267–79. doi:10.1016/j.cell.2016.04.028 PubMed PMID: 27180905.

50. Chaung K, Baharav TZ, Henderson G, Zheludev IN, Wang PL, Salzman J. SPLASH: A statistical, reference-free genomic algorithm unifies biological discovery. Cell. 2023 Dec 7;186(25):5440-5456.e26. doi:10.1016/j.cell.2023.10.028 PubMed PMID: 38065078.

51. Sharma E, Sterne-Weiler T, O’Hanlon D, Blencowe BJ. Global Mapping of Human RNA-RNA Interactions. Mol Cell. 2016 May 19;62(4):618–26. doi:10.1016/j.molcel.2016.04.030 PubMed PMID: 27184080.

52. Ziv O, Gabryelska MM, Lun ATL, Gebert LFR, Sheu-Gruttadauria J, Meredith LW, et al. COMRADES determines in vivo RNA structures and interactions. Nat Methods. 2018 Oct;15(10):785–8. doi:10.1038/s41592-018-0121-0 PubMed PMID: 30202058; PubMed Central PMCID: PMC6168409.

53. Sugimoto Y, Vigilante A, Darbo E, Zirra A, Militti C, D’Ambrogio A, et al. hiCLIP reveals the in vivo atlas of mRNA secondary structures recognized by Staufen 1. Nature. 2015 Mar 26;519(7544):491– 4. doi:10.1038/nature14280 PubMed PMID: 25799984; PubMed Central PMCID: PMC4376666.

54. Cai Z, Cao C, Ji L, Ye R, Wang D, Xia C, et al. RIC-seq for global in situ profiling of RNA–RNA spatial interactions. Nature. 2020 Jun 18;582(7812):432–7. doi:10.1038/s41586-020-2249-1

55. Gay LA, Turner PC, Renne R. Modified Cross-Linking, Ligation, and Sequencing of Hybrids (qCLASH) to Identify MicroRNA Targets. Curr Protoc. 2021 Oct;1(10):e257. doi:10.1002/cpz1.257 PubMed PMID: 34610213; PubMed Central PMCID: PMC8500481.

56. Wu T, Cheng AY, Zhang Y, Xu J, Wu J, Wen L, et al. KARR-seq reveals cellular higher-order RNA structures and RNA–RNA interactions. Nat Biotechnol. 2024 Dec;42(12):1909–20. doi:10.1038/s41587-023-02109-8

57. Ye R, Hu N, Cao C, Su R, Xu S, Yang C, et al. Capture RIC-seq reveals positional rules of PTBP1-associated RNA loops in splicing regulation. Mol Cell. 2023 Apr 20;83(8):1311-1327.e7. doi:10.1016/j.molcel.2023.03.001 PubMed PMID: 36958328.

58. Helwak A, Tollervey D. Mapping the miRNA interactome by crosslinking ligation and sequencing of hybrids (CLASH). Nat Protoc. 2014 Mar;9(3):711–28. doi:10.1038/nprot.2014.043 PubMed PMID: 24577361; PubMed Central PMCID: PMC4033841.

59. Lee WH, Zhang M, Lu Z. Integrated Analysis of Cross-Link-Ligation Data for the Detection of RNA 2D/3D Structures and Interactions In Vivo. In: Hamada M, Fukunaga T, Zeng C, editors. RNA-RNA and RNA-DNA Interactions [Internet]. New York, NY: Springer US; 2026 [cited 2026 Apr 10]. p. 53– 77. Available from: https://doi.org/10.1007/978-1-0716-4670-0_3 doi:10.1007/978-1-0716-4670-0_3

60. Hsi-Yang Fritz M, Leinonen R, Cochrane G, Birney E. Efficient storage of high throughput DNA sequencing data using reference-based compression. Genome Res. 2011 May;21(5):734–40. doi:10.1101/gr.114819.110 PubMed PMID: 21245279; PubMed Central PMCID: PMC3083090.

61. Thorvaldsdóttir H, Robinson JT, Mesirov JP. Integrative Genomics Viewer (IGV): high-performance genomics data visualization and exploration. Brief Bioinform. 2013 Mar;14(2):178–92. doi:10.1093/bib/bbs017 PubMed PMID: 22517427; PubMed Central PMCID: PMC3603213.

62. Zhang M, Li K, Bai J, Van Damme R, Zhang W, Alba M, et al. A snoRNA–tRNA modification network governs codon-biased cellular states. Proc Natl Acad Sci U S A. 120(41):e2312126120. doi:10.1073/pnas.2312126120 PubMed PMID: 37792516; PubMed Central PMCID: PMC10576143.

63. Kiss T. Small nucleolar RNA-guided post-transcriptional modification of cellular RNAs. EMBO J. 2001 Jul 16;20(14):3617–22. doi:10.1093/emboj/20.14.3617

64. Tycowski KT, Smith CM, Shu MD, Steitz JA. A small nucleolar RNA requirement for site-specific ribose methylation of rRNA in Xenopus. Proc Natl Acad Sci. 1996 Dec 10;93(25):14480–5. doi:10.1073/pnas.93.25.14480

65. Babraham Bioinformatics - FastQC A Quality Control tool for High Throughput Sequence Data [Internet]. [cited 2025 Apr 30]. Available from: https://www.bioinformatics.babraham.ac.uk/projects/fastqc/

66. Bolger AM, Lohse M, Usadel B. Trimmomatic: a flexible trimmer for Illumina sequence data. Bioinformatics. 2014 Aug 1;30(15):2114–20. doi:10.1093/bioinformatics/btu170

67. Dobin A, Davis CA, Schlesinger F, Drenkow J, Zaleski C, Jha S, et al. STAR: ultrafast universal RNA-seq aligner. Bioinforma Oxf Engl. 2013 Jan 1;29(1):15–21. doi:10.1093/bioinformatics/bts635 PubMed PMID: 23104886; PubMed Central PMCID: PMC3530905.

68. McInnes L, Healy J, Melville J. UMAP: Uniform Manifold Approximation and Projection for Dimension Reduction [Internet]. arXiv; 2020 [cited 2025 Jul 1]. Available from: http://arxiv.org/abs/1802.03426 doi:10.48550/arXiv.1802.03426

69. Robinson MD, Oshlack A. A scaling normalization method for differential expression analysis of RNA-seq data. Genome Biol. 2010 Mar 2;11(3):R25. doi:10.1186/gb-2010-11-3-r25

70. Wu Y, Shao W, Yan M, Wang Y, Xu P, Huang G, et al. Transfer learning enables identification of multiple types of RNA modifications using nanopore direct RNA sequencing. Nat Commun. 2024 May 14;15(1):4049. doi:10.1038/s41467-024-48437-4

71. Li Y, Zhang C, Feng C, Pearce R, Lydia Freddolino P, Zhang Y. Integrating end-to-end learning with deep geometrical potentials for ab initio RNA structure prediction. Nat Commun. 2023 Sep 16;14(1):5745. doi:10.1038/s41467-023-41303-9

72. Sato K, Akiyama M, Sakakibara Y. RNA secondary structure prediction using deep learning with thermodynamic integration. Nat Commun. 2021 Feb 11;12(1):941. doi:10.1038/s41467-021-21194-4

73. Wang Y, Pan Z, Mou M, Xia W, Zhang H, Zhang H, et al. A task-specific encoding algorithm for RNAs and RNA-associated interactions based on convolutional autoencoder. Nucleic Acids Res. 2023 Nov 27;51(21):e110. doi:10.1093/nar/gkad929

